# Metabolic complementarity between a brown alga and associated cultivable bacteria provide indications of beneficial interactions

**DOI:** 10.1101/813683

**Authors:** Bertille Burgunter-Delamare, Hetty KleinJan, Clémence Frioux, Enora Fremy, Margot Wagner, Erwan Corre, Alicia Le Salver, Cédric Leroux, Catherine Leblanc, Catherine Boyen, Anne Siegel, Simon M. Dittami

**Author notes:** CEBEDEAU, Research and Expertise Center for Water, Allée de la découverte, 11 (B53), Quartier Polytech 1, B-4000, Liège, Belgium (Hetty KleinJan). These authors contributed equally to this study (Bertille Burgunter-Delamare, Hetty KleinJan).

## Abstract

Brown algae are key components of marine ecosystems and live in association with bacteria that are essential for their growth and development. *Ectocarpus siliculosus* is a genetic and genomic model for brown algae. Here we use this model to start disentangling the complex interactions that may occur between the algal host and its associated bacteria. We report the genome-sequencing of 10 alga-associated bacteria and the genome-based reconstruction of their metabolic networks. The predicted metabolic capacities were then used to identify metabolic complementarities between the algal host and the bacteria, highlighting a range of potentially beneficial metabolite exchanges between them. These putative exchanges allowed us to predict consortia consisting of a subset of these ten bacteria that would best complement the algal metabolism. Finally, co-culture experiments were set up with a subset of these consortia to monitor algal growth as well as the presence of key algal metabolites. Although we did not fully control but only modify bacterial communities in our experiments, our data demonstrated a significant increase in algal growth in cultures inoculated with the selected consortia. In several cases, we also detected, in algal extracts, the presence of key metabolites predicted to become producible via an exchange of metabolites between the alga and the microbiome. Thus, although further methodological developments will be necessary to better control and understand microbial interactions in *Ectocarpus*, our data suggest that metabolic complementarity is a good indicator of beneficial metabolite exchanges in holobiont.

## 1 Introduction

Microbial symbionts are omnipresent and important for the development and functioning of multicellular eukaryotes. Together the eukaryote hosts and their microbiota form meta-organisms also called holobionts. Elucidating the interactions within microbial communities and how they affect host physiology is a complex task and requires an understanding of the dynamics within the microbiome and the host, as well as of possible inter-species interactions and/or metabolic exchanges that could occur between the partners. One way to dissect those interactions is via targeted co-culture experiments using culturable bacteria. This approach works particularly well for 1:1 or 1:2 interactions, but as the number of potentially interacting organisms increases, selecting the “right” bacterial consortia becomes a major bottleneck (Lindemann *et al.* 2016).

Metabolic complementarity has previously been proposed as an indicator for potentially beneficial host-symbiont interactions and can be assessed *in silico* using the metabolic networks of the host and the microbiota (Dittami, Eveillard, *et al.* 2014; Levy *et al.* 2015). Common examples of metabolic complementarity are associations of autotrophic and heterotrophic organisms such as corals and their photosynthetic symbionts (Rohwer *et al.* 2002), or algae, and their heterotrophic bacterial biofilm (Wahl *et al.* 2012). In this case, the autotrophic partner has a metabolic capacity (photosynthesis) that allows for the production of metabolic intermediates (organic carbon), which can be further metabolized by the heterotrophic partners. However, especially in systems with long-lasting interactions more complex metabolic interdependencies are likely to evolve (*e.g.* Amin *et al.* 2015).

As a tool to further explore such interactions, Frioux et al. (Frioux *et al.* 2018) have proposed the pipeline MiSCoTo. Given the metabolic networks of a host and several symbionts, this tool predicts potential metabolic capacities of one partner that could be unlocked by a contribution of a metabolite from another (e.g. the provision of carbohydrates by a photosynthetic organism unlocking the biochemical processes related to primary metabolism in heterotrophs). Furthermore, this computational approach uses these complementarities to define minimal consortia (i.e. with the lowest possible number of exchanges/contributors) allowing the host to reach its maximum metabolic potential. However, the actual predictive value of these models, both in terms of the effect on host growth and fitness, and in terms of the metabolic scope (i.e. the metabolites producible by the holobiont system), remains to be assessed.

Here we have applied the MiSCoTo tool to the filamentous brown alga *Ectocarpus siliculosus*, a model filamentous brown alga with an available metabolic network (Prigent *et al.* 2014), as well as a selection of 10 *Ectocarpus*-derived bacteria (KleinJan *et al.* 2017). We then selected specific minimal microbial consortia for *in vivo* testing of the proposed hypotheses (growth rate, production of specific metabolites). Our results demonstrate a clear positive effect of inoculation with the predicted bacterial consortia on algal growth as well as an effect on the production of algal metabolites predicted to depend on bacterial contributions. *In vivo* observations largely corresponded to *in silico* predictions despite the incomplete input data (with models limited to annotated pathways) and the fact that we had only limited control of the microbiome. The present work thus generates numerous testable hypotheses on specific beneficial interactions between *Ectocarpus* and its microbiome, but also provides a proof of concept for the overall predictive power of network-based metabolic complementarity for beneficial host-microbe interactions.

## 2 Methods

### 2.1 Bacterial cultures and genome sequencing

Ten bacterial strains were selected from the 46 isolated by KleinJan et al. from *Ectocarpus subulatus* (KleinJan *et al.* 2017). They were grown in liquid Zobell and/or diluted R2A until bacterial growth was visible with the naked eye (∼3 days at room temperature), and their identity was confirmed by sequencing of the 16S rRNA gene with the primers 8F and 1492R (KleinJan *et al.* 2017). Bacterial DNA was extracted using the UltraClean® Microbial DNA isolation kit (MoBio, Qiagen, Hilden, Germany) and used for standard pair-end sequencing at the GENOMER platform (FR2424, Station Biologique de Roscoff), using Illumina Miseq technology (V3 chemistry, 2×300bp). After cleaning with Trimmomatic v0.38, default parameters (Bolger *et al.* 2014), the paired-end reads were assembled using SPADES v3.7.0 (Bankevich *et al.* 2012; default parameters for long reads). The RAST/SEED server (Aziz *et al.* 2008) was used for gene annotation, and sequences were later also incorporated into the MAGE platform (Vallenet *et al.* 2006).

### 2.2 *In silico* predictions of metabolic interactions and selection of consortia

Bacterial metabolic networks were constructed using Pathway Tools version 20.5 (Karp *et al.* 2016) and version 2 of the *Ectocarpus siliculosus* EC32 metabolic network for the host, prior to any gap-filling step, in order to prevent the presence of possibly false positive reactions in the model. (because these false positive reactions could hide algal bacterial interactions). This network comprised a total of 2,118 metabolites, 1,887 metabolic reactions, and was able to produce five of the 50 metabolites known to be a part of the *Ectocarpus* biomass (Aite *et al.* 2018) with only the culture medium as input. For the remaining 45 compounds the lack of producibility can be explained by the presence of metabolic gaps – either because a reaction was missed during the reconstruction of the network (missing annotation etc.), or because the corresponding pathways require metabolite exchanges with other partners in the environment, *e.g.* bacteria. The more such gaps can be filled by exchanging compounds between two metabolic networks, the higher we consider the degree of metabolic complementarity between the corresponding organisms.

Here we used the MiSCoTo tool (Frioux *et al.* 2018) to compute such potential metabolic exchanges between *Ectocarpus* and any of the ten targeted bacteria. The underlying model of MiSCoTo assumes that a compound is producible by a host-symbiont community if there is a chain of metabolic reactions which transforms the culture medium into the expected compound without taking into consideration flux accumulations or competition for resources, and allowing for the exchange of compounds across cell boundaries. These simplifications imply that compounds predicted to be producible *in silico* may, in some cases, remain unproducible *in vivo*, although the consortium has all the genes to activate the pathways.

In this study MiSCoTo was run twice, first to determine the scope of all algal compounds that become producible via exchanges with all 10 bacterial genomes together, and as second time to select minimal bacterial consortia for the production of these compounds. In both cases the Provasoli culture medium was used as a source as defined previously (Prigent *et al.* 2014).

### 2.3 Algal cultures

Two of the six predicted bacterial consortia were tested experimentally via algal-bacterial co-culture experiments. Additionally, each member of the two consortia was tested individually, as well one other sequenced strain that was not part of any of the predicted minimal consortia, *i.e. Sphingomonas* sp. 391. *Ectocarpus siliculosus* (strain 32; accession CCAP 1310/4, origin San Juan de Marcona, Peru) was cultured under standard conditions (13 °C; 12h light regime) in Provasoli-enriched natural seawater until the start of antibiotic treatment. Prior to co-culture experiments, algal filaments were treated with a mixture of the following liquid antibiotics: 45 μg/ml Penicillin G, 22.5 μg/ml streptomycin, and 4.5 μg/ml chloramphenicol dissolved in Provasoli-enriched artificial seawater 450 mM Na^+^, 532 mM Cl^-^, 10 mM K^+^, 6 mM Ca^2+^, 46 mM Mg^2+^, 16 mM SO_4_ ^2-^. Filaments were exposed to 25 ml of this solution for 3 days and then placed in Provasoli-enriched artificial seawater for 3 days to recover. The absence of bacteria on the algal surface was verified by microscopy using phase-contrast (Olympus BX60, 1.3-PH3 immersion objective, 800x magnification) and by plating of algal filaments on Petri dishes with Zobell medium followed by three weeks of incubation at room temperature.

### 2.4 Co-culture experiments

For co-culture experiments, cell densities of bacterial cultures were determined using a BD FACS CantoTM II flow cytometer (BD Bioscience, San Jose, CA) using samples fixed in Tris-EDTA. Before the start of the experiment, antibiotic-treated algae (three replicate cultures per condition) were inoculated with 2.3*10^5^ bacterial cells per strain and ml medium. Each co-culture was then incubated for 4 weeks under standard algal growth conditions (see above). During this time, algal growth was quantified by measuring the filament length of the algae each week using the binocular microscope (3 measurements per replicate). Furthermore, bacterial abundance in the algal growth medium was estimated using flow cytometry (described above) and bacteria attached to algal cell walls were counted by microscopy (5× 10 μm long filaments observed per biological replicate, 800x magnification in phase contrast). At the end of the experiment, general algal morphology was observed using a LEICA DMi8 microscope and in parallel, remaining algal tissues were frozen in liquid nitrogen and freeze-dried for downstream analyses. Two controls (three replicates each) were run in parallel: a non-antibiotic treated positive control (CTRL w/o. ATB), and an antibiotic-treated non-inoculated alga as a negative control (CTRL w. ATB).

### 2.5 Bacterial community composition after co-culture experiments

A metabarcoding approach was implemented to investigate the composition of the bacterial community after the co-culture experiments. For each culture, 20 mg ground freeze-dried tissue (TissueLyserII Qiagen, Hilden, Germany; 2×45sec, 30 Hz) was used for DNA extraction (DNeasy Plant Mini Kit, Qiagen; standard protocol). Nucleotide concentrations were verified with NanodropONE (Thermofisher Scientific). A mock community comprised of DNA from 32 bacterial strains (covering a variety of taxa) as well as a negative control were included in addition to the samples (see Thomas *et al.* in prep. for details). Libraries were prepared according to the standard Illumina protocol for metabarcoding MiSeq technology targeting the V3–V4 region (Illumina 2017) and sequenced using Illumina MiSeq Technology (2×300 bp, pair-end reads; MiSeq Reagent v3 kit; Platform de Séquencage-Génotypage GENOMER, FR2424, Roscoff).

Resulting raw sequences (7,354,164 read pairs) were trimmed using fastq_quality_trimmer from the FASTX Toolkit (quality threshold 30; minimum read length 200) and assembled into 6,804,772 contigs using PandaSeq v2.11 (Masella *et al.* 2012). Data were analyzed with Mothur (V.1.40.3) according to the MiSeq Standard Operating Procedures (Kozich *et al.* 2013). Contigs were pre-clustered (allowing for four mismatches), and aligned to the Silva_SEED 132b database for sequence classification. Chimeric sequences were removed (Vsearch) and the remaining sequences classified taxonomically (Wang *et al.* 2007). Non-bacterial sequences were removed and the remaining sequences were then clustered into operational taxonomic units (OTUs) at a 97% identity level and each OTU was classified to the genus level where passible (Wang *et al.* 2007). All OTUs with n ≤ 10 sequences were removed resulting in a final data matrix with 1,834,992 sequences. The OTU matrix was subsampled to have the same number of sequences per sample for downstream analyses.

### 2.6 Targeted metabolomics

Seven metabolites predicted to be producible by the algae only in presence of metabolic exchanges with specific bacteria were selected for targeted metabolite profiling after manual verification of automatic predictions of corresponding pathways in the algal and bacterial networks and based on their biological importance for the alga: L-histidine, putrescine, beta-alanine, nicotinic acid, folic acid, auxin, and spermidine. Metabolites were extracted from 10 mg of ground, freeze-dried tissue using a triple extraction protocol based on the method of Bligh and Dyer (1959): two ml of methanol:chloroform:water (6:4:1) were used as first extraction solvent, then the remaining pellet was extracted with 1 ml of chloroform:methanol (1:1), and finally, a 3rd extraction was performed using 1ml of H_2_O. The supernatants of each extraction were pooled and evaporated under a stream of nitrogen. The residue was then resuspended in 100 μl methanol:water (1:1) and analyzed on an ACQUITY Ultra-performance convergence chromatography (UPC^2^) system (Waters®, Milford, USA) equipped with a Viridis BEH column (3×100 mm, 1.7 μm). A linear gradient of two solvents was used to separate peaks: supercritical carbon dioxide (Solvent A), and methanol spiked with 0.1% formic acid (Solvent B). The gradient ran from 5% to 25% of solvent B (35% for spermidine and nicotinic acid) during 2 minutes, was kept at this level for another 2 minutes and then gradually reduced back to 5% during 3 minutes. The UPC^2^ system was coupled to a Xevo G2 Q-Tof mass spectrometer (Waters), operating in positive ESI ion mode (m/z 20–500). Blanks, as well as standards of all 7 compounds obtained from Sigma-Aldrich (St. Louis, MO, USA), were run in parallel to samples. The resulting chromatograms were then used to examine the presence/absence of the target compounds in the other samples based on retention time and the mass spectra. Analyses were performed at the METABOMER platform (FR2424, Station Biologique de Roscoff).

### 2.7 Statistical analyses

Growth data (both algal and bacterial) were confirmed to follow a normal distribution using a Shapiro-Wilk test (Rstudio v1.0.44). Significant differences between all treatments after four weeks of co-culture (day 28) were calculated with an ANOVA and a Tukey honestly significant difference (HSD) post-hoc test with a significance level α 0.05 using the PAST software version 3.20 (Hammer *et al.* 2001).

## 3 Results

### 3.1 Predicted metabolic interactions and selection of beneficial bacterial consortia

Genome sequencing and subsequent bioinformatics analyses yielded bacterial genome assemblies with sufficient coverage and 11-72 scaffolds per genome Table 1). Metabolic networks were then reconstructed for these ten genomes. On average, 1,714 reactions, 111 transport reactions, and 1,405 metabolites (Table 2) were predicted per bacterium. These reactions belonged, again on average, to 261 pathways, 137 of which were complete and 124 were incomplete (*i.e.* missing one or more reactions). Based on metabolic complementarity analysis carried out using MiSCoTo, these bacterial networks were predicted to enable the production of 160 additional compounds with the algal networks, including several polyamines (Cadaverine, Spermidine, Agmatine), amino acids (Histidine, Tyrosine, beta-alanine), vitamins B3, B9, and E, several lipids and lipid derivatives, and nucleic acids. Please refer to Supplementary Table S1 for a complete list of compounds. Many of these compounds were also previously predicted via the metabolic interaction between the same strain of *E. siliculosus* and the associated bacterium *Candidatus* Phaeomarinobacter ectocarpi (Dittami, Barbeyron, *et al.* 2014; Prigent *et al.* 2017). A total of six bacterial consortia comprising three bacterial strains each (Table 3) were predicted to be sufficient to enable the production of all of these compounds. Of these six proposed consortia, two comprising one phylogenetically distinct bacterium each (i.e. the *Bacteriodetes Imperialibacter* vs the *Gammaproteobacterium Marinobacter*) were chosen for *in vivo* testing using algal-bacterial co-cultures.

**Table 1:**
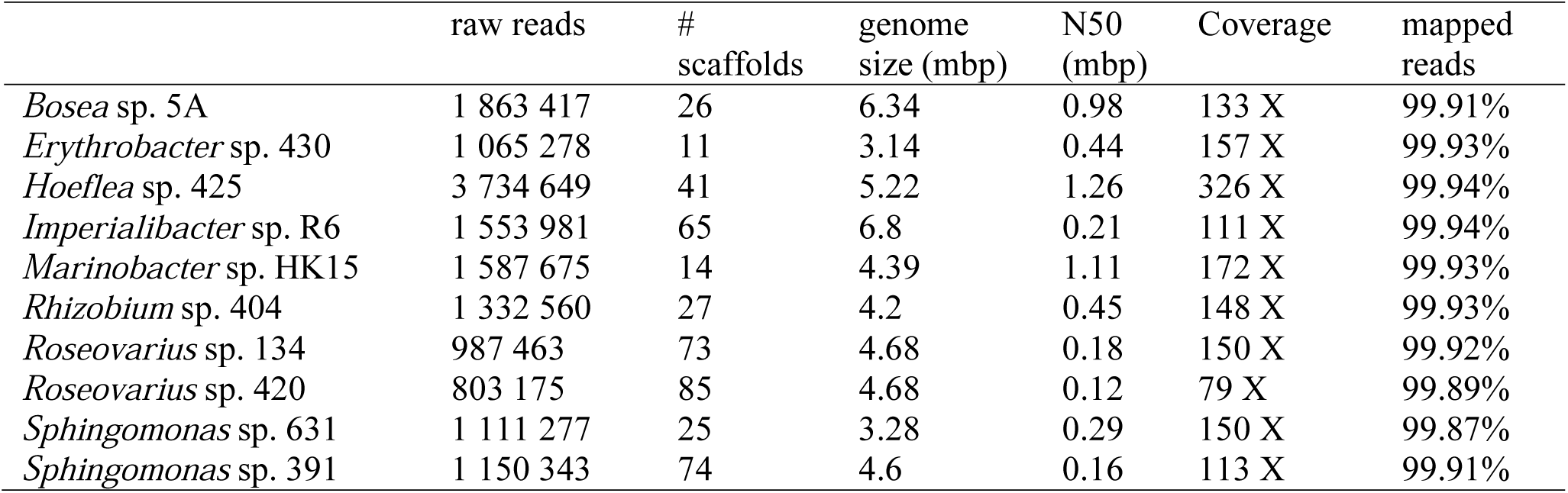
Overview of bacterial genomes used in this study and corresponding assembly statistics.

**Table 2:** Predicted metabolic pathways (complete pathways in parentheses), reactions and metabolites in bacterial metabolic networks.

|  | number of<br>pathways | number of<br>reactions | transport<br>reactions | number of<br>metabolites |
| --- | --- | --- | --- | --- |
| *Bosea* sp. 5A | 298 (187) | 1892 | 153 | 1557 |
| *Erythrobacter* sp. 430 | 218 (91) | 1532 | 63 | 1247 |
| *Hoeflea* sp. 425 | 315 (170) | 1920 | 129 | 1558 |
| *Imperialibacter* sp. R6 | 239 (131) | 1711 | 100 | 1425 |
| *Marinobacter* sp. HK15 | 249 (128) | 1679 | 128 | 1364 |
| *Rhizobium* sp. 404 | 289 (142) | 1814 | 125 | 1462 |
| *Roseovarius* sp. 134 | 263 (146) | 1703 | 125 | 1418 |
| *Roseovarius* sp. 420 | 263 (143) | 1701 | 125 | 1418 |
| *Sphingomonas* sp. 361 | 224 (108) | 1519 | 69 | 1239 |
| *Sphingomonas* sp. 391 | 254 (126) | 1671 | 92 | 1358 |

**Table 3:** Minimal bacterial consortia predicted by MiSCoTo that enabled the production of 160 algal compounds. See Supporting table S1 for a detailed list of compounds.

| Solution proposed by MiSCoTo | In vivo testing? |
| --- | --- |
| *Marinobacter* sp. HK15, *Roseovarius* sp. 420, *Hoeflea* sp. 425 | Yes |
| *Roseovarius* sp. 420, *Imperialibacter* sp. R6, *Hoeflea* sp. 425 | Yes |
| *Marinobacter* sp. HK15, *Bosea* sp. 5a, *Roseovarius* sp. 420 | No |
| *Marinobacter* HK15, *Hoeflea* sp. 425, *Roseovarius* sp. 134 | No |
| *Imperialibacter* sp. R6, *Hoeflea* sp. 425, *Roseovarius* sp.134 | No |
| *Marinobacter* sp. HK15, *Bosea* sp. 5a, *Roseovarius* sp. 134 | No |

### 3.2 Growth rates in co-culture experiments

The inoculation with one or several bacterial strains significantly enhanced algal growth by a factor of 2 compared to controls (Figure 1A). This positive effect was observed both for the predicted bacterial consortia and for all the individual strains tested. At the same time, the abundance of bacteria on algal filaments after four weeks of cultivation was significantly lower in cultures initially inoculated with bacteria compared to both controls with and without initial antibiotic treatment (Figure 1B), although bacterial cell counts in the medium were similar between co-culture experiments and the non-inoculated control after 28 days (Supplementary Figure S1).

**Figure 1:**
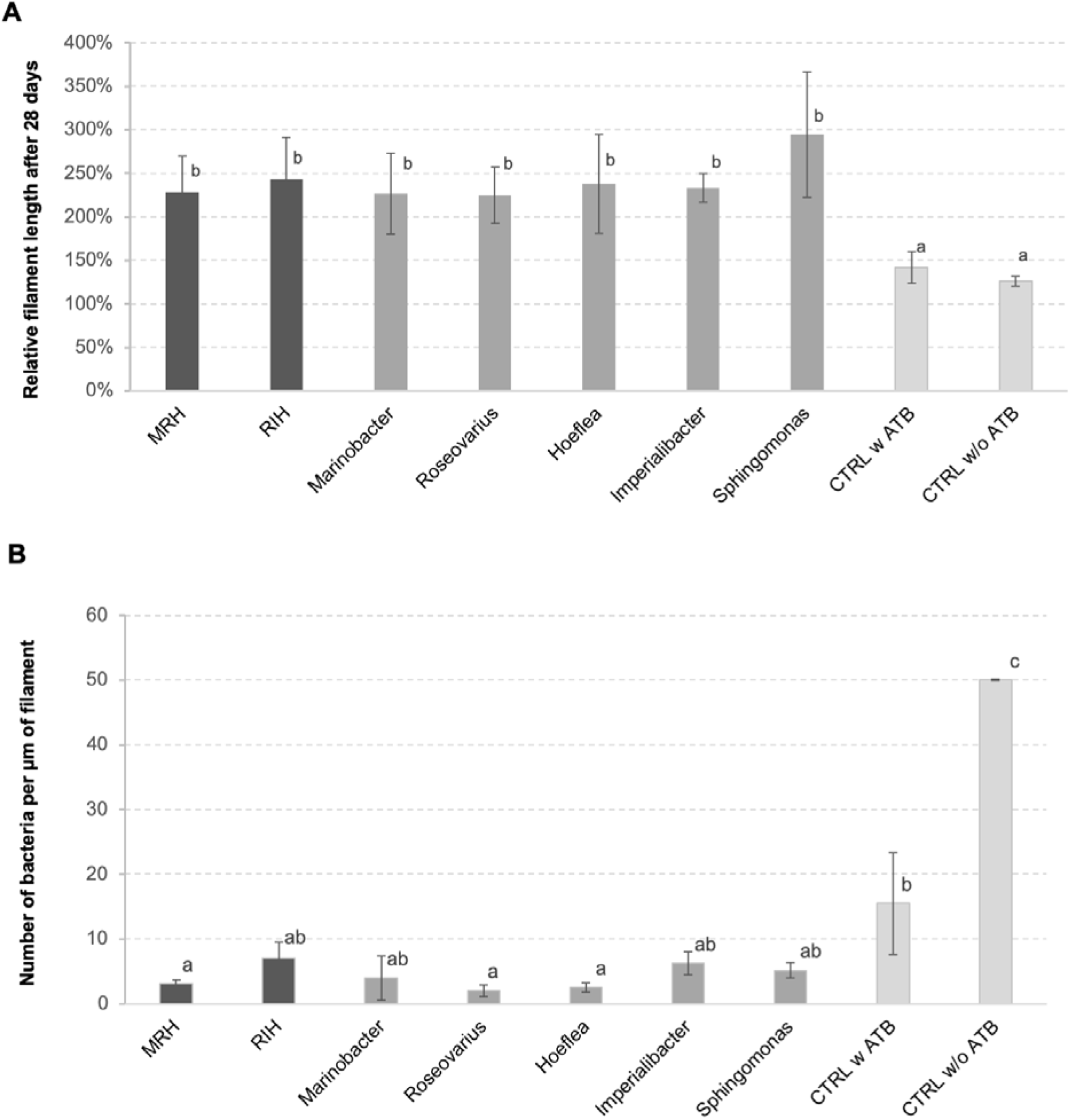
A) Relative length of *E. siliculosus* filaments after 28 days of (co-)culture compared to the starting point. B) Number of bacteria detected on algal filaments after 28 days of co-culture. Both panels A and B show means of 3 replicate co-cultures ± SD and differences are statistically significant (one-way ANOVA p<0.01). The letters above the columns indicate the results of a TUKEY HSD pairwise comparisons (p<0.05). CTRL = control, ATB = antibiotic treatment, MRH = *Marinobacter*-*Roseovarius*-*Hoeflea*; RIH = *Roseovarius*-*Imperialibacter*-*Hoeflea*.

### 3.3 Bacterial impact on morphology

Compared to the negative control, which exhibited a ball-like morphology typical for “axenic” cultures (Tapia *et al.* 2016), all bacterial inocula tested resulted in filamentous thalli with clear branching patterns (Figure 2). We furthermore observed differences in the branching patterns depending on the bacterial inocula. For example, *Sphingomonas*-inoculated cultures produced relatively long filaments with few branching sites (Figure 2H), whereas *Hoeflea*-inoculated cultures produced filaments with frequent branching (Figure 2E). *Imperialibacter* induced aggregation of individual filaments (Figure 2F), while in all other co-cultures, filaments remained more or less separated. These differences were, however, difficult to quantify given complexity of their morphology.

**Figure 2:**
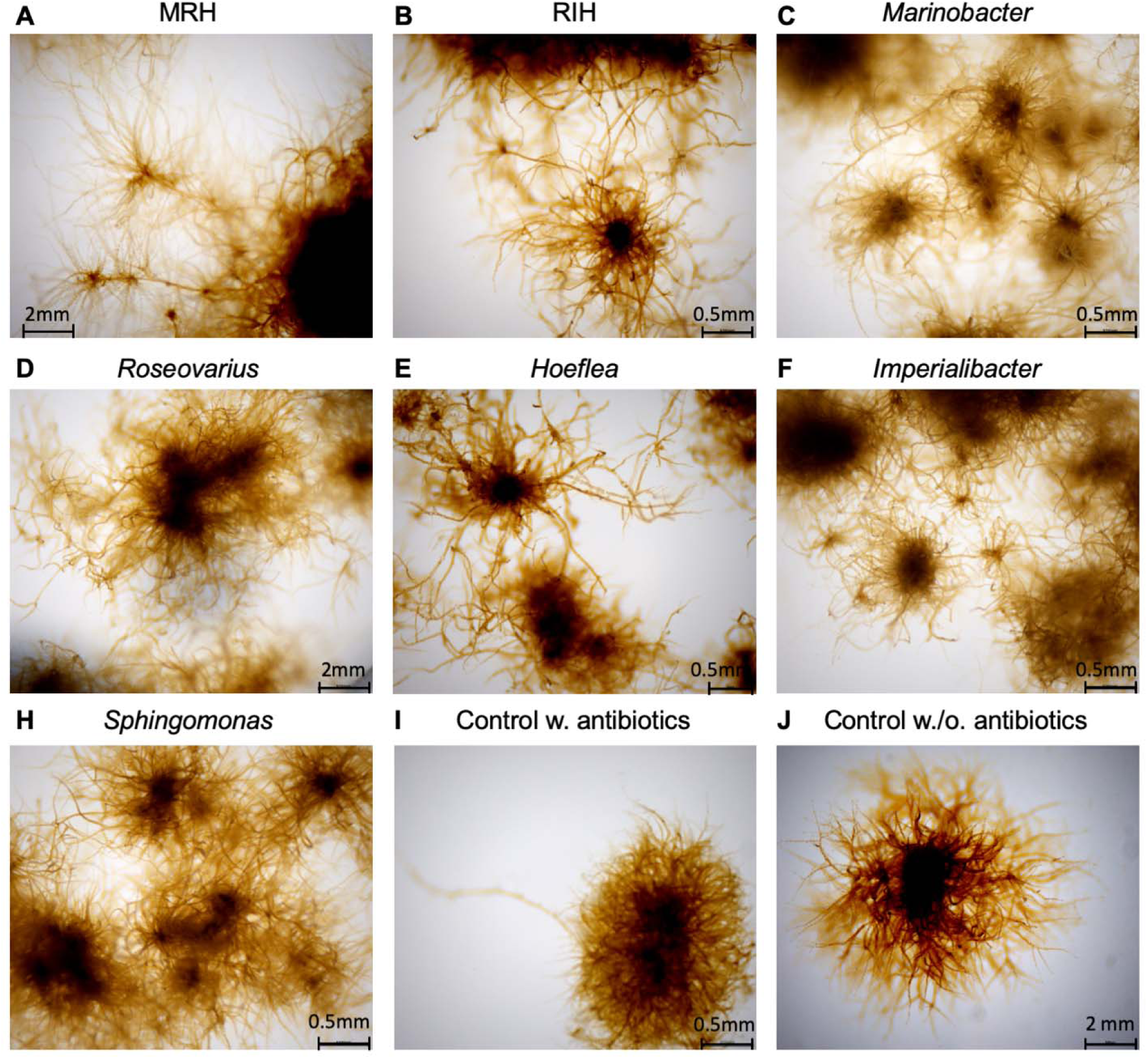
Morphological effect co-cultures with bacteria on *E. siliculosus* after 4 weeks of co-culturing. MRH = *Marinobacter*-*Roseovarius*-*Hoeflea*, RIH = *Roseovarius*-*Imperialibacter*-*Hoeflea*.

### 3.4 (Algal) metabolome in co-culture conditions

Seven putatively key metabolites (l-histidine, putrescine, beta-alanine, nicotinic acid, folic acid, auxin, and spermidine) predicted to be non-producible by the alga alone but producible via exchanges with some bacterial consortia, were quantified in algal tissues by UPC^2^-MS after four weeks of co-culture. The presence/absence of these metabolites is shown in Figure 3, comparing both the predicted producibility by metabolic network analysis and the experimental UPC^2^-MS results. In the negative control, i.e. antibiotic-treated algae that were not inoculated with bacteria, none of the compounds could be identified by UPC^2^-MS confirming the computational predictions. In contrast, in all co-cultures, at least one target compound was experimentally detected. Furthermore, each compound became producible in at least one of the co-cultures. Overall, across the 56 predictions made based on the metabolic networks (7 metabolites x 8 consortia including the individual bacteria and the negative control) *in silico* and *in vivo* data agreed in 28 cases (Figure 3). Only in four cases did we observe the presence of a metabolite although it was not predicted by the networks. Finally, in 24 cases we did not detect the presence of a metabolite predicted to be producible in the co-cultures.

**Figure 3:**
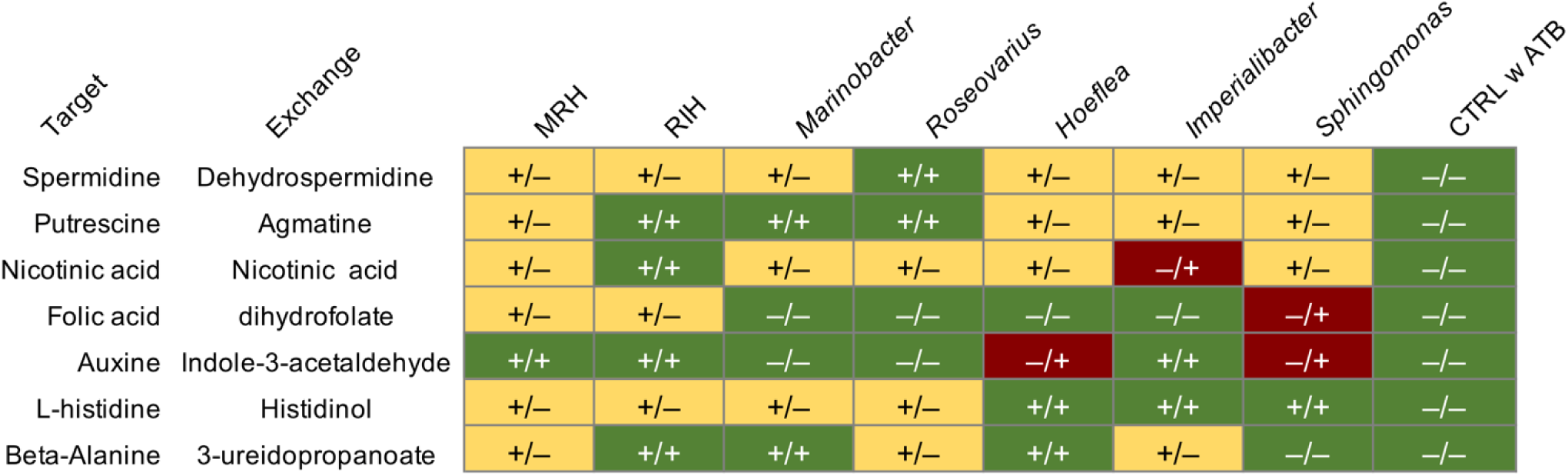
Comparison of predicted production of target metabolites in co-cultures based on metabolic networks (symbol before the slash) and results from targeted UPC^2^-MS analyses of algal filaments after 28 days (symbol after the slash). The column “Exchange” indicates one possible compound provided by the microbiome leading to the production of the compound in the column “Target” in the algal metabolome; it was these target metabolites that were tested for using UPC^2^-MS. All experiments were carried out in triplicate, each replicate of the same condition yielding identical results. (-): a target metabolite was not predicted/detected (+): a metabolite was predicted/detected. Green highlights conditions where predictions correspond to the in vivo observations, red highlights compounds that were detected although no pathway was predicted. Yellow indicates compounds potentially producible via bacterial exchanges that were not detected. MRH: *Marinobacter*-*Roseovarius*-*Hoeflea*; RIH: *Roseovarius*-*Imperialibacter*-*Hoeflea*; CTRL = control; ATB = antibiotic treatment.

### 3.5 Bacterial community composition after co-culture experiments

The bacterial community composition of each sample was analyzed by 16S rDNA metabarcoding at the end of the co-culture experiments. This was done to verify if the bacteria inoculated had grown in the co-cultures and to determine to what extent other bacteria were present and affected by the inoculations. The results (Table 4) show that, except for *Imperialibacter*, all of the bacterial strains inoculated were detected in the corresponding co-cultures 28 days after inoculation. However, except for *Marinobacter* and *Hoeflea*, read abundances of these strains were low compared to the total number of reads. In parallel, several other OTUs that had not been inoculated were detected in our co-culture experiments, suggesting that these bacteria were at least partially resistant to or protected from (*e.g.* within the cell wall) the antibiotic treatments applied, and were able to recover under the experimental conditions: in total 30 additional OTUs with a minimal abundance of 1% of total reads were detected in our samples, accounting for 63 to 82% of the total reads. Furthermore, *Hoeflea* reads were dominant in all samples including the non *Hoeflea*-inoculated cultures (14-30% of total reads).

**Table 4:** Observed abundance of target OTUs after four weeks of co-culture. The table shows number of reads obtained corresponding to each OTU (mean three replicates ± SD). Bold numbers on grey background indicate OTUs expected to be present based on the inoculations.

|  | MRH | RIH | *Marino-*<br>*bacter* | *Roseo-*<br>*varius* | *Hoef-*<br>*lea* | *Im-*<br>*periali-*<br>*bacter* | *Sphing-*<br>*omonas* | CTRL<br>w. ATB | CTRL<br>w/o.<br>ATB |
| --- | --- | --- | --- | --- | --- | --- | --- | --- | --- |
| *Marinobct.O*<br>TU00030 | **82 ± 48** | 0 ± 0 | **1103 ± 1068** | 0 ± 0 | 0 ± 0 | 0 ± 0 | 38 ± 38 | 0 ± 0 | 1 ± 1 |
| *Roseovarius*<br>OTU00055 | **8 ± 7** | **11 ± 3** | 1 ± 1 | **41 ± 10** | 1 ± 1 | 0 ± 0 | 0 ± 0 | 0 ± 0 | 0 ± 0 |
| *Hoeflea*<br>OTU00001 | **10265 ± 1586** | **7644 ± 889** | 4483 ± 2777 | 15635 ± 1349 | **15321 ± 3515** | 13426 ± 5338 | 10216 ± 4345 | 8899 ± 2811 | 3618 ± 1055 |
| *Imperialib.*<br>OTU00044 | 0 ± 0 | **0 ± 0** | 0 ± 0 | 0 ± 0 | 0 ± 0 | **0 ± 0** | 0 ± 0 | 0 ± 0 | 0 ± 0 |
| *Sphingomn.*<br>OTU00097 | 0 ± 0 | 0 ± 0 | 0 ± 0 | 0 ± 0 | 0 ± 0 | 1 ± 1 | **4 ± 4** | 0 ± 0 | 0 ± 0 |
| Other OTUs* | 39403 ± 2138 | 23458 ± 1828 | 26223 ± 3187 | 36374 ± 7810 | 34190 ± 5508 | 38076 ± 4292 | 29066 ± 3302 | 42323 ± 9670 | 28009 ± 5897 |

\* see Supplementary Figure S2 for details

## 4 Discussion

### Metabolic complementarity, a powerful metric despite limitations

Metabolic complementarity intuitively seems like an excellent marker for beneficial metabolic interactions. The more organisms are complementary at the metabolic and by extension the gene level, the more they can potentially benefit from each other (Levy *et al.* 2015); the more they overlap in terms of metabolic pathways, the more likely they are to compete for the same resources (Kreimer *et al.* 2012). There are, however, two important restrictions that limit the applicability of this simple idea. First, the possibility of a beneficial exchange does not necessarily mean that it will occur, because this may require the presence and activation of excretion/uptake mechanisms in both partners, *e.g.* via chemical or environmental cues. Secondly, the genome-scale metabolic models used to predict metabolic complementarities may be partially erroneous and incomplete. For instance, metabolic networks frequently do not comprise interactions of chemical signals with receptor molecules, which may be key to regulate interactions (Zhou *et al.* 2016; Wang *et al.* 2018). Furthermore, in many cases, they are based on automatic predictions and annotations of protein sequences, which may, in some cases, miss genes or introduce overpredictions of functions (Schnoes *et al.* 2009). In this paper, we provide first *in vivo* tests of host-microbe interactions inferred from genome-based predictions of metabolic complementarity. Despite the aforementioned restrictions and simplifications, our results discussed below provide a strong indication that, genome-based predictions of metabolic complementarity is a powerful tool to handle the complexity of host microbe systems and to generate hypotheses on their interactions.

### Similar complementarities found across studies and *Ectocarpus* symbionts

Compared to a previous analysis of metabolic complementarity between *Ectocarpus* and another associated bacterium, *Candidatus* Phaeomarinobacter ectocarpii, (Dittami, Barbeyron, *et al.* 2014; Prigent *et al.* 2017), newly producible compounds predicted in this study were largely similar, notably regarding polyamines, histidine, beta-alanine, and auxin. This similarity persists even though metabolic complementarity analyses were performed using MiSCoTo, which incorporates the notion of different compartments minimizing the number metabolite exchanges (Frioux *et al.* 2018) and despite the fact that different bacteria were examined. The main difference compared to the previous study is that numerous additional compounds were predicted to be exchanged, which can be explained by the fact that ten rather than one bacterial network were available to complete the algal network.

### Inoculation with metabolically complementary bacteria enhances growth rate and impacts morphology and metabolism

As described above, both the bacterial consortia tested, as well as all of the bacteria inoculated individually had clear positive effects on algal growth and impacted algal morphology and metabolite profiles, even though, by the time the co-cultures were harvested, some of the inoculated bacteria were present only in very low abundance or even below the detection limit. These positive effects could be due either to interactions early in the co-culture experiments followed by a decline in bacterial abundance, or due to the capacity of bacteria to impact and interact with their algal hosts even at very low cell concentrations. The latter would support the hypothesis that part of the observed effects may not be due to the exchanges of (abundant) primary metabolites, such as the predicted histidine/histidinol, but due to lowly concentrated signaling molecules or growth hormones. One such compound could be the examined auxin, which was detected in 5 of the 7 tested co-cultures, and which has previously been shown to modify the developmental patterns and morphology of *Ectocarpus* cultures (Le Bail *et al.* 2010) in a similar way as bacterial inoculations. Another observation was that the abundance of bacteria on algal filaments but not in the medium was significantly lower in co-culture conditions compared to the controls. This suggests that the inoculated bacteria, either directly, or indirectly, by stimulating algal growth or defense, can also regulate biofilm formation (see Goecke *et al.* 2010 for a review).

Interestingly, although differences in the effects of individual bacteria and bacterial consortia were observed on metabolite profiles and morphology, all consortia had similar effects on algal growth. Indeed, all of the tested bacteria, including *Sphingomonas*, which was not part of the minimal solutions proposed by MiSCoTo, were to a great extent complementary to the alga, already covering a large part of the metabolic gaps. In future studies, it may be particularly useful to incorporate a larger range of bacteria, possibly from other sources so that they are not expected to have evolved mutualistic interactions with brown algae. These negative controls could then be used to correlate growth rates with the presence or absence of specific metabolic capacities in the network. Once the list of candidate metabolite exchanges has been narrowed down by such comparisons, supplying these metabolites from artificial sources but also testing for their excretion into the medium by bacteria can be used to corroborate their role.

### Predicted metabolic exchanges likely to occur in part

With respect to the predictions of target metabolites, we observed that for a large number of cases, predictions from the metabolic networks corresponded to the observations made by experimental metabolic profiling: none of the target metabolites were detected in the negative control, and only in four cases (Figure 3), did we detect compounds in co-cultures that were not predicted to be there. This could either be attributed to undetected metabolic pathways in the examined/added bacteria (*e.g.* due to missing annotations) or, more likely, to the activity of other bacteria present in our co-culture experiments (see below). Furthermore, there were several cases in which a potentially co-producible metabolite was not detected in our co-cultures. Here two explanations appear particularly likely: first, the metabolites in question may be produced but quickly metabolized in certain consortia, so that they do not accumulate sufficiently to be detectable in our cultures; secondly, it is possible that the corresponding biosynthetic pathway of the metabolite was not active or that the necessary exchange of metabolites was not taking place. To resolve this point in future experiments, the addition of gene expression data may help to establish whether or not biosynthetic or degradation pathways are active. From a global perspective, however, the fact that none of the compounds in question were detected in negative controls, but all of them it at least one co-culture condition, constitutes a highly promising result.

### Outlook

In our opinion, the main challenge for future *in vivo* studies of metabolic complementarity will be to better control the *Ectocarpus*-associated microbiome in co-culture experiments, and thus to avoid any impact of non-inoculated microbes. The currently applied antibiotic treatments are successful in removing bacteria from the algal surface to a level where they are no longer detectable by microscopy and spreading on culture medium, but once the treatment is stopped and algae are left to recover, so do parts of the microbiome, possibly from spores that were inactive or embedded in the algal cell wall and thus less susceptible to our treatments (Tetz and Tetz 2017). In the light of these results, we strongly recommend routine metabarcoding analysis for any type of coculture experiment, also in other model systems. One possibility in the future would be to use axenization protocols based on the movement of gametes, as has been done for *Ulva mutablilis* (Spoerner *et al.* 2012); at least some strains of *Ectocarpus* have previously been shown to produce phototactic gametes (Kawai *et al.* 1990). A second alternative is the continuous use of antibiotics throughout the experiment, and working with antibiotic-resistant bacterial strains. In this context a better understanding of the metabolic requirements of the algae will help to durably maintain axenic cultures.

Despite these challenges, the present study constitutes an important proof of concept for the use of metabolic complementarity to study simplified system of mutualistic host-symbiont interactions. We anticipate that, in the long run, this concept can be applied not only to controlled co-culture experiments, but that it will also prove useful for the interpretation of more complex datasets such as metatranscriptomic or metagenomic data.

## 6 Author Contributions

Conceived the experiments: BB, HK, SD; Conceived in silico analyses: CF, AS. Performed experiments: BB, HK; Performed analyses: BB, ALS, EF, CF, MW, SD, EC, CL; Wrote the manuscript: SD, HK, BB. Corrected and approved of the final manuscript: all authors.

## 7 Funding

This work was funded partially by the CNRS Momentum call, the ANR project IDEALG (ANR-10-BTBR-04) “Investissements d’Avenir, Biotechnologies-Bioressources”, the European Union’s Horizon 2020 research and innovation Programme under the Marie Sklodowska-Curie grant agreement number 624575 (ALFF), and joint PhD scholarship from the Brittany region (Project HOSALA) and the Sorbonne University (ED227).

### 8 Acknowledgments

We thank Laurence Dartevelle for advice on algal culturing; Sylvie Rousvoal for extractions of bacterial DNA; Gwenn Tanguy and Erwan Legeay from the GENOMER platform (FR2424, Station Biologique de Roscoff) for access to the sequencing platform and support during library preparation; and François Thomas, Angélique Gobet, Maeva Brunet for helpful discussions and for sharing their mock community; Dominique Marie and Christian Jeanthon for granting us access to their flow cytometer; and the ABIMS platform and the GenOuest bioinformatics core facility (https://www.genouest.org) for providing the computing infrastructure for amplicon analyses and community predictions. The LABGeM (CEA/Genoscope & CNRS UMR8030), the France Génomique and French Bioinformatics Institute national infrastructures (funded as part of Investissement d’Avenir program managed by Agence Nationale pour la Recherche, contracts ANR-10-INBS-09 and ANR-11-INBS-0013) are acknowledged for support within the MicroScope annotation platform.

## 11 Supplementary Material

**Supplementary Table S1**: Metabolites predicted to become producible by the alga as a result of metabolite exchanges between the alga and bacteria. (uploaded separately)

**Supplementary Figure S1:**
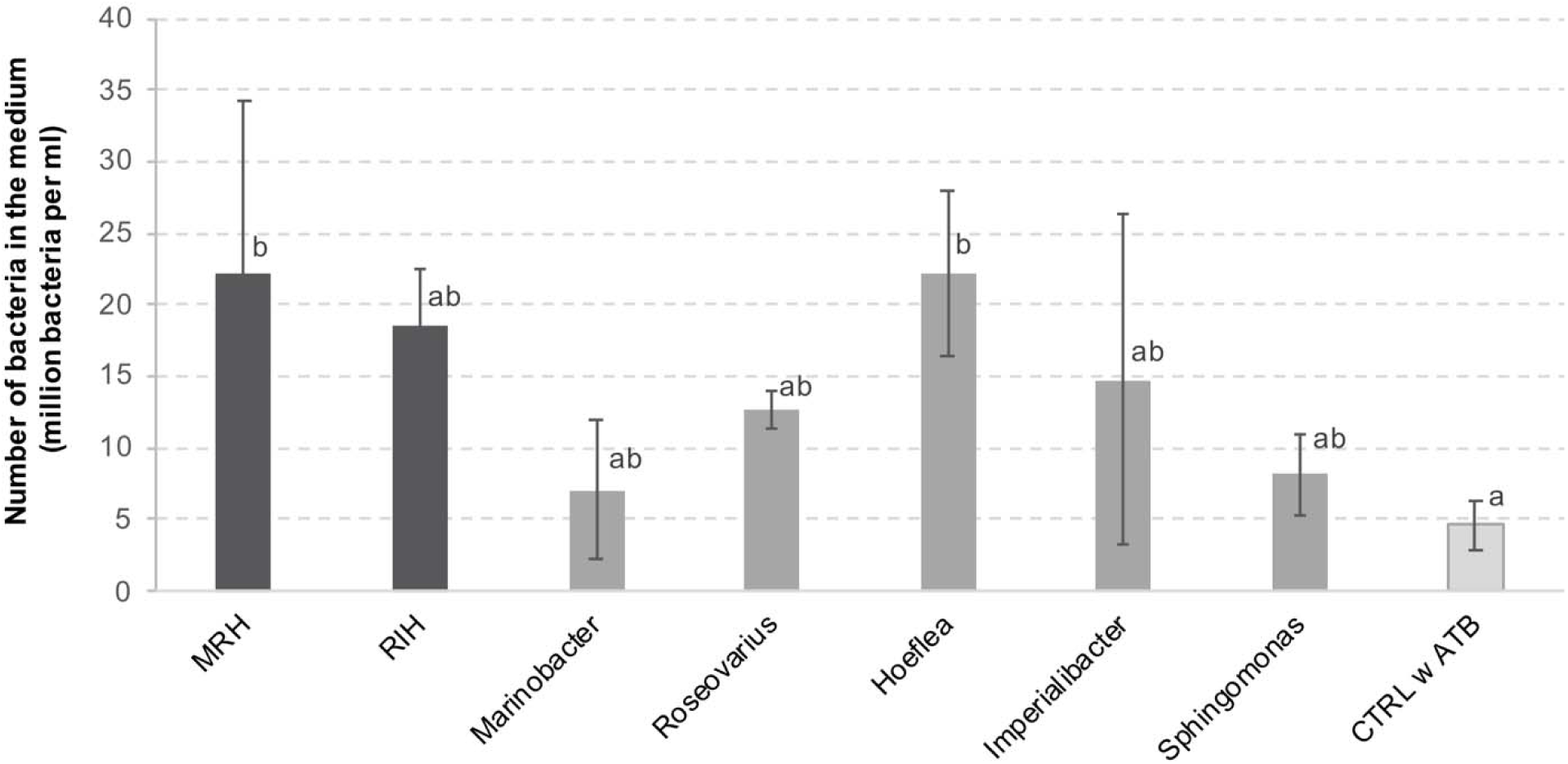
Number of bacteria detected in the algal culture medium after 28 days of co-culture. The graph shows means of 3 replicates ± SD and differences are statistically significant (one-way ANOVA p<0.01). The letters above the columns indicate the results of a TUKEY HSD pairwise comparisons (p<0.05). CTRL = control, ATB = antibiotic treatment.

**Supplementary Figure S2:**
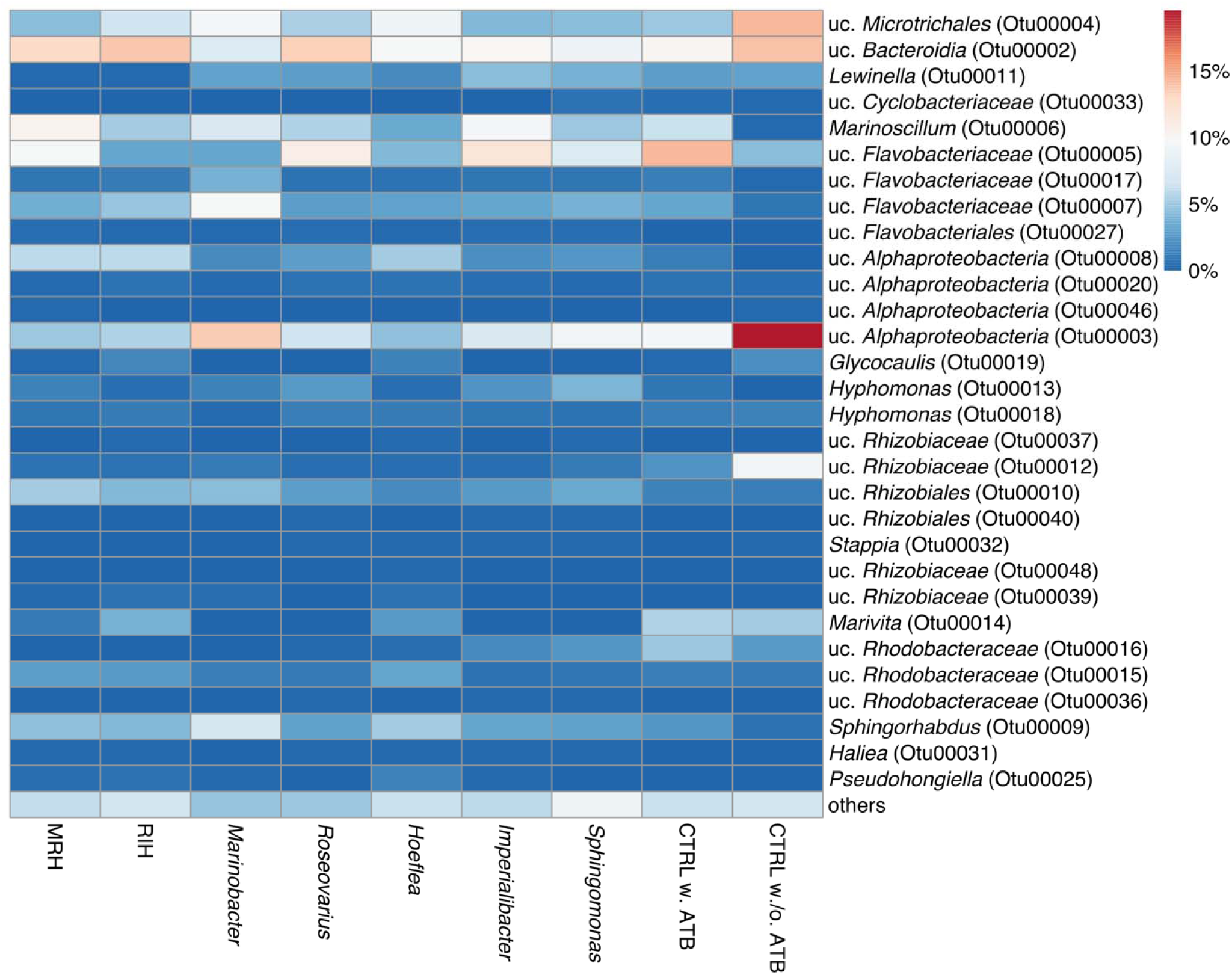
Heatmap of relative OTU abundance for all 30 OTUs that made up over 1% of the total number of reads and that were not inoculated (See Table 4 for the latter). This heatmap as generated using the ClustVis service (Metsalu and Vilo 2015) using “correlation” as a distance measure and “average linkage” as clustering method. The color code corresponds to the mean sequence abundance for each OTU in the three replicates a percentage of total reads; uc. = unclassified

## 12 Data Availability Statement

The metabarcoding data generated for this study has been deposited at the European Nucleotide Archive (ENA) under project accession number PRJEB34356. The bacterial genomes have been deposited at the ENA under the sample accessions ERZ1079053-ERZ1079062.

